# An MRI Atrophy Biomarker Predicts Global Prognosis in Early *De Novo* Parkinson’s Disease

**DOI:** 10.1101/528810

**Authors:** Yashar Zeighami, Seyed-Mohammad Fereshtehnejad, Mahsa Dadar, D. Louis Collins, Ronald B. Postuma, Alain Dagher

## Abstract

**Background:** Commonly used neuroimaging biomarkers in Parkinson’s disease (PD) are useful for diagnosis but poor at predicting outcomes. We explored whether an atrophy pattern from whole-brain structural MRI, measured in the drug-naïve early stage, could predict PD prognosis.

**Methods:** 362 *de novo* PD patients with T1-weighted MRI (n=222 for the main analysis, 140 for the validation analysis) were recruited from the Parkinson’s Progression Markers Initiative (PPMI). We investigated a previously identified PD-specific network atrophy pattern as a potential biomarker of disease severity and prognosis. Progression trajectories of motor function (MDS-UPDRS-part III), cognition (Montreal Cognitive Assessment (MoCA)), and a global composite outcome measure were compared between atrophy tertiles using mixed effect models. The prognostic value of the MRI atrophy measure was compared with ^123^I ioflupane single photon emission computed tomography, the postural-instability-gait-disturbance score, and cerebrospinal fluid markers.

**Findings:** After 4.5 years follow-up, PD-specific atrophy network score at baseline significantly predicted change in UPDRS-part III (r=-0.197, p=0.003), MoCA (r=0.253, p=0.0002) and global composite outcome (r=-0.249, p=0.0002). Compared with the 3^rd^ tertile (i.e. least atrophy), the tertile with the highest baseline atrophy (i.e. the 1st tertile) had a 3-point annual faster progression in UPDRS-part III (p=0.012), faster worsening of posture-instability gait scores (+0.21 further annual increase, p<0.0001), faster decline in MoCA (−0.74 further annual decline in MoCA, *p*=0.0372) and a +0.38 (*p*=0.0029) faster annual increase in the global composite z-score. All findings were replicated in a validation analysis using 1.5T MRI. By comparison, the other biomarkers were limited in their ability to predict prognosis either in the main or validation analysis.

**Interpretation:** A PD-specific network atrophy pattern predicts progression of motor, cognitive, and global outcome in PD, and is a stronger predictor of prognosis than any of the other tested biomarkers. Therefore, it has considerable potential as a prognostic biomarker for clinical trials of early PD.

## INTRODUCTION

Parkinson’s disease (PD) is a complex neurodegenerative disorder with a broad range of motor and non-motor features. It is notably heterogeneous, varying considerably in its clinical manifestations and prognosis.^1^ This represents an important confound in clinical trials against PD. Therefore, prognostic biological markers are urgently needed to stratify and monitor patients for clinical trials. Most described markers of prognosis are based on clinical measures (e.g. the postural instability and gait disturbance (PIGD) score)^2^, while biomarkers including cerebrospinal fluid (CSF) measures, gene mutations, and nigrostriatal dopamine tracers have not performed well as predictors of prognosis.^3^

Neuroimaging techniques have particular potential as prognostic markers, since they directly measure brain morphological and functional changes. Positron emission tomography (PET) and single-photon emission CT (SPECT) (e.g. with the DAT tracer ^123^| ioflupane) can directly identify nigrostriatal neurodegeneration (even years before the motor signs of PD appear),^3^ qualifying them as valid diagnostic biomarkers. Nevertheless, while they have been validated as diagnostic tests, it remains unclear to what degree dopaminergic imaging techniques can predict PD progression.

Because of magnetic resonance imaging (MRI)’s broad availability and standardized acquisition parameters, any MRI biomarker of prognosis would have potential for widespread application. Although routine clinical MRI scans are classically considered as normal in PD, recent methodological improvements may make MRI prognostic markers feasible.4 However, there is currently no accepted MRI prognostic biomarker for PD.5 Using high resolution 3T MRI data from the Parkinson’s Progression Markers Initiative (PPMI),6 we previously introduced an MRI-based whole-brain atrophy measure that was strongly associated with disease severity at baseline.7 In the current study, we investigated whether this PD-network atrophy pattern could predict the rate of progression of motor and non-motor symptoms. Second, we compared the predictive power of the MRI-based biomarker with other potential predictors, including motor examination, postural/gait measures, CSF measures, and SPECT. Third, we measured the rate of change of the MRI biomarker in longitudinal MRI scans from PPMI. Finally, we replicated our analysis in a separate validation set of the PPMI population scanned with 1.5T MRI.

## METHODS

### Participants

PD patients with age ≥30, a diagnosis of PD within the last 2 years, a baseline Hoehn and Yahr stage of I or II, and no anticipated need for symptomatic treatment within six months of entry were recruited by the Parkinson’s Progression Markers Initiative (PPMI) (see: http://www.ppmi-info.org), a multicenter international study of de novo individuals with early idiopathic PD.6 Selection criteria included asymmetric resting tremor or asymmetric bradykinesia or two of bradykinesia, resting tremor and rigidity, plus confirmation of a dopaminergic deficit using dopamine transporter imaging (see below). For the current study, we also excluded any participant with less than one year of follow-up or no MRI available. Therefore, 362 de novo treatment-naive PD patients were included. Of these, 222 had 3T MRI and 140 had 1.5T MRI.

The relevant local institutional review boards approved the PPMI protocol and written informed consent was obtained from all participants prior to inclusion. We retrieved data from the PPMI database in October 2017 in compliance with the PPMI data use agreement. The average follow-up period was 4.5 years.

### Brain Imaging Analysis

High resolution T1-weighted 3T MRI was available in 232 PD as well as 117 healthy age-matched control (HC) participants at baseline. Deformation-based morphometry (DBM) was used as a measure of brain atrophy. The analysis was performed as explained previously.^7^ Each participant’s MRI was first linearly and then nonlinearly registered to the Montreal Neurological Institute (MNI) ICBM-152 template.^8^ The nonlinear transformations were used to calculate the Jacobian determinant of the deformation matrix at each voxel for each participant, to yield individual DBM or atrophy maps. We then performed independent component analysis (ICA) on the DBM maps^9^ to identify PD-specific atrophy distribution in early PD (**Figure 1**).

**Figure 1.**
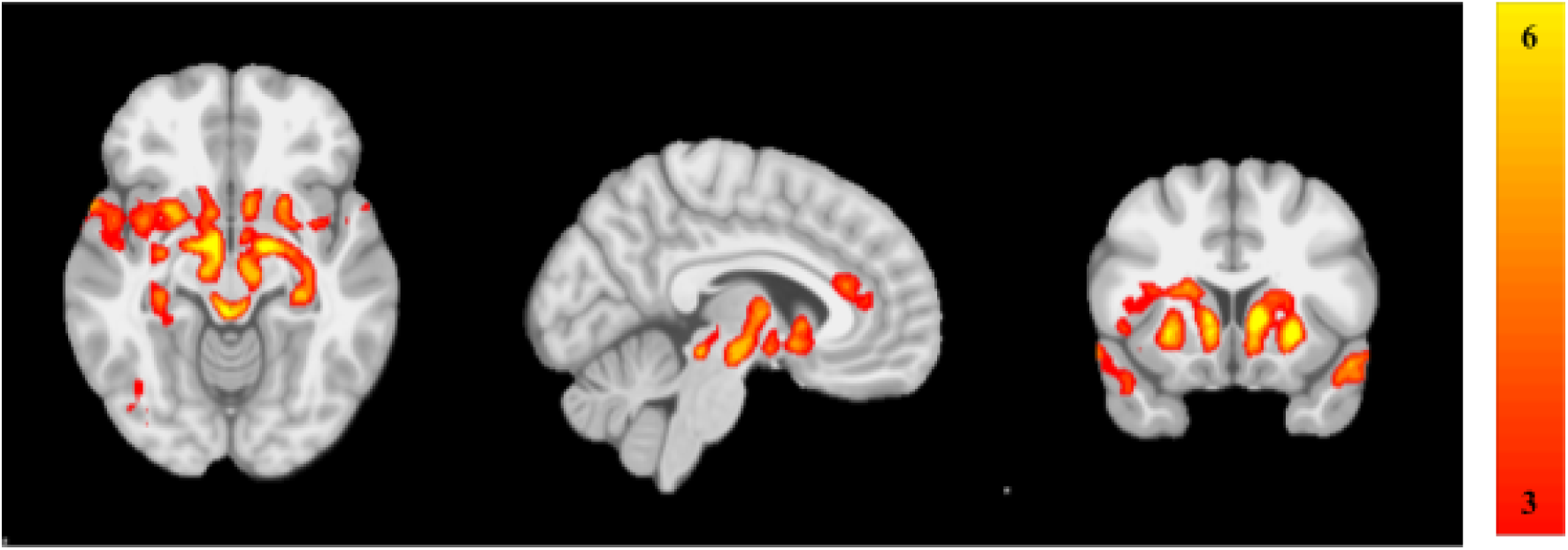
Deformation-based morphometry (DBM) maps of the Parkinson’s disease. (PD)-specific network showing significant differences in atrophy between PD individuals and healthy controls (p=0.003 after Bonferroni correction for multiple comparison). The independent component analysis (ICA) spatial map was converted to a z-statistic image via a normalized mixture–model fit and then thresholded at z = 3. Selected sections in Montreal Neurological Institute (MNI) space at coordinates Z= −10, X= −6, Y= +14) (From: *Zeighami Y et al eLife 2015*)

The PD-related atrophy score was calculated as a single numerical indicator of atrophy for each participant, and is referred to here as the PD-network atrophy score. The atrophy scores were transformed into z-scores (lower values represent more severe atrophy). For the longitudinal analysis of brain morphometry, all T1-weighted MRIs for subjects with up to 4-years follow up (average of 2.2 years follow up) were included. This data includes 222 PD patients at baseline, 142 at year-1, 127 at year-2, and 84 at year-4 (Subjects were not scanned at year-3 based on data downloaded in May 2018). Healthy control subjects included 112 subjects at baseline, 64 at year-1, and 20 at year-4. The drop from 232 PD to 222 and 117 HC to 112 is due to lack of follow up information in other domains needed for statistical analysis. All procedures were repeated as explained for baseline analysis.

Furthermore, to validate our results in an out-of-sample set of participants, we computed individual DBM maps in a validation set of 140 different PPMI PD participants who had undergone T1-weighted MRI at 1.5T.^7^ The atrophy pattern identified in the 3T sample^7^ was then applied to the DBM images of the 1.5T data to calculate the atrophy score within this validation sample. To do so, for each individual in the 1.5T independent sample, we multiplied the DBM value at each voxel by the PD-network atrophy score from the same voxel. We averaged these values to obtain a weighted atrophy score for each individual. Due to lack of follow up for subjects with 1.5T T1-weighted MRI imaging, longitudinal neuroimaging analyses are limited to the main dataset.

### Baseline and Follow-up Clinical Assessments

Demographic characteristics including age, sex, race, family history, duration of symptoms, and education level were recorded at the screening visit. A comprehensive set of clinical features including both motor and non-motor signs and symptoms was evaluated at baseline and each follow-up visit. Motor-related measures consist of Movement Disorder Society-Unified Parkinson’s Disease Rating Scale (MDS-UPDRS)-Part II, MDS-UPDRS-Part III, PIGD score,^2^ and Schwab and England activities of daily living (ADL) score. The non-motor symptoms and signs used here are non-motor experiences of daily living (MDS-UPDRS-Part I), autonomic dysfunction [the Scales for Outcomes in PD-Autonomic (SCOPA-AUT) total score],^10^ depression [Geriatric Depression Scale (GDS) score],^11^ anxiety [State-Trait Anxiety Inventory (STAI) score],^12^ REM sleep behaviour disorder (RBD) [RBD screening questionnaire (RBDSQ) score],^13^ sleep disturbances [Epworth Sleepiness Score (ESS)]^14^ and impulse control disorders (ICD) [Questionnaire for Impulsive-Compulsive Disorders in Parkinson’s disease (QUIP) score]^15^ and global cognition [age and education adjusted Montreal Cognitive Assessment (MoCA) score].^16^

We also created a global composite outcome (GCO) score as a single numeric indicator of overall disease severity, as previously described.^17,18^ This was standardized by combining the z-scores of the most clinically relevant motor and non-motor manifestations of disease progression, namely non-motor symptoms (MDS-UPDRS-Part I), motor symptoms (MDS-UPDRS-Part II), motor signs (MDS-UPDRS-Part III), overall activities of daily living (Schwab and England ADL), and global cognitive function (MoCA). For calculating follow-up z-scores we applied the mean and standard deviation (SD) from baseline as the reference state to measure progression over time. The GCO was calculated by averaging these z-scores at baseline and at the most recent follow-up visit for each participant. Higher GCO scores indicate worse overall function.^18^

### Clinical Subtypes

We recently recommended a guideline for subtyping *de novo* PD participants based on clinical features,^18^ in which three distinct subtypes of PD were defined (’mild motor predominant’, ‘diffuse malignant’, and ‘intermediate’) based on the combination of a composite motor severity score, RBD, dysautonomia, and cognitive impairment. To investigate the ability of the PD-network atrophy to predict a change in clinical subtype category, we recalculated the subtype assignments after 4.5 years follow-up using the updated reference percentile values derived from the score distributions of these motor and non-motor features in the entire PPMI population. Furthermore, we defined tremor-dominant, intermediate and PIGD motor phenotypes (as described previously)^2^ at baseline and investigated the association between baseline PD-network atrophy score and the shift in motor phenotypes after follow-up.

### Other Potential Predictive Biomarkers

Cerebrospinal fluid (CSF) was collected for all participants at baseline. CSF amyloid-beta (A*β*1-42), total Tau (T-tau), and phosphorylated tau (P-tau181) were measured by INNO-BIA AlzBio3 immunoassay (*Innogenetics* Inc.) and *α*-synuclein was measured by enzyme-linked immunosorbent assay.^19^ We calculated the CSF A*β*42/T-tau ratio as recommended in a recent publication.^20^

SPECT with the DAT tracer ^123^| ioflupane was obtained in 351 participants with PD both at baseline and after two years.^6^ Using the occipital lobe as a reference region, striatal binding ratio (SBR) was calculated for the left and right caudate and putamen separately in each individual.

### Statistical Methods

All relevant data available through the PPMI website were used. Missing values (<4%) were imputed by using the mean value for the entire cohort. Univariate correlation between the baseline PD-network atrophy score and clinical measures was investigated using *Pearson* correlation. We used *Bonferroni* correction to adjust for type I error inflation induced by multiple comparisons. Multivariate linear regression was performed to regress out the effect of age on the associations between PD-network atrophy score and clinical measures. We applied receiver operating characteristic (ROC) curve analysis to compare the accuracy of various biomarkers at baseline to predict a 1.5 SD increase in GCO after 4.5 years of follow-up. The threshold of 1.5 SD (the average increase in GCO=0.8 SD) was chosen to include a reasonable number of participants who experienced a more rapid (almost double) progression rate (n=38 out of 222). Area under the curve (AUC) and its 95% confidence interval (CI) were calculated for each biomarker.

The population was divided into tertiles of PD-network atrophy score -and other biomarkers-to test the a-priori hypothesis that participants with mild, moderate and severe atrophy at baseline would progress differently over time. We chose to divide the cohort into three for this purpose based on our previous hierarchical clustering analysis of this dataset.^18^ We then applied mixed effect models to compare the longitudinal trajectories of clinical outcomes and their interaction with age between the three subgroups defined by the baseline level of the biomarker of interest. We used a separate model for each clinical outcome (i.e. UPDRS-III, MoCA, GCO, and PIGD score). In each model we included clinical outcome, age, their interaction, as well as sex and education as fixed effects and subject as a random effect (equation 1)

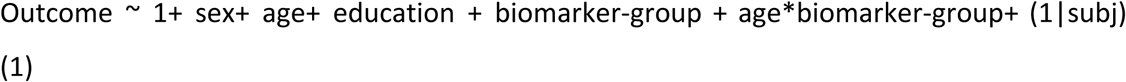

For the longitudinal brain imaging analysis, the PD atrophy score was calculated for all PD and HC subjects for each visit. We then used a mixed effect model to investigate the interaction between aging and disease status (Cohort = PD or HC).

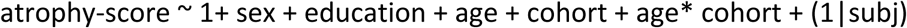

Furthermore to investigate the effect of baseline atrophy in the PD group on the longitudinal changes of the atrophy score, we used the same model to investigate other longitudinal outcomes with baseline tertiles within the PD cohort.

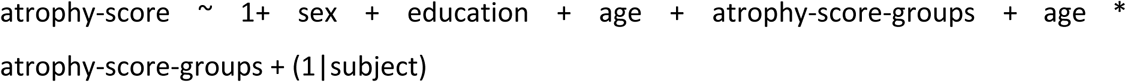

All models were implemented in MATLAB 2015b using the fitlme function. Other univariate and multivariate analyses were performed using *IBM SPSS Statistics* software (version 23.0 and *R* version 3.2.2. A two-tailed *p*-value of <0.05 was considered as the threshold for statistically significant differences or associations in all analyses unless otherwise specified.

## RESULTS

3T MRI data were available for 232 individuals of the PPMI population at baseline, of whom 222 subjects had at least one year of follow-up data. **Table 1** summarizes demographic and clinical characteristics of the study population including both main and validation cohorts at baseline and after 4.5 years.

**Table 1.**
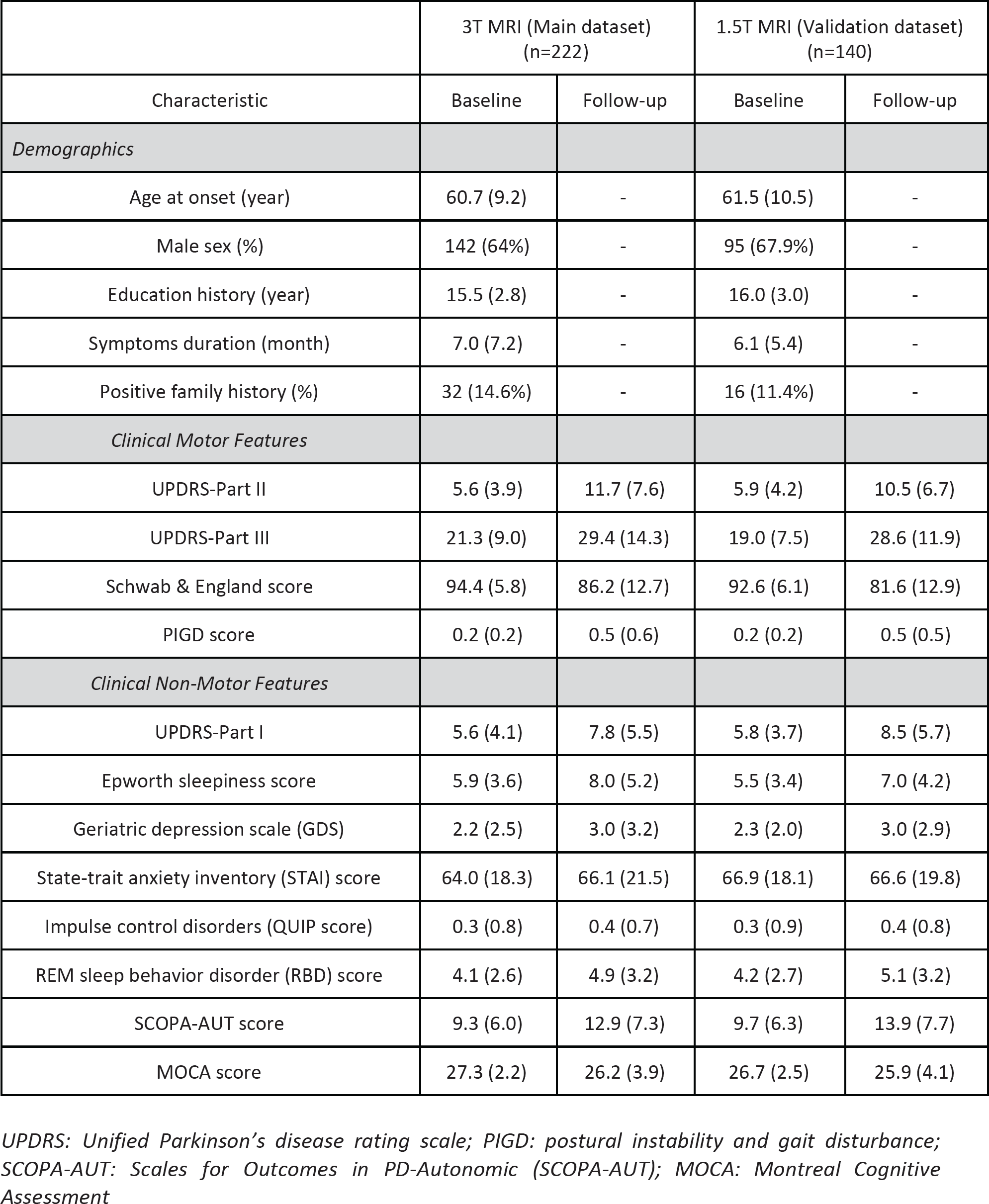
Demographic and clinical characteristics of the study population

|  | 3T MRI (Main dataset)<br>(n=222) |  | 1.5T MRI (Validation dataset)<br>(n=140) |  |
| --- | --- | --- | --- | --- |
| Characteristic | Baseline | Follow-up | Baseline | Follow-up |
| *Demographics* |  |  |  |  |
| Age at onset (year) | 60.7 (9.2) | - | 61.5 (10.5) | - |
| Male sex (%) | 142 (64%) | - | 95 (67.9%) | - |
| Education history (year) | 15.5 (2.8) | - | 16.0 (3.0) | - |
| Symptoms duration (month) | 7.0 (7.2) | - | 6.1 (5.4) | - |
| Positive family history (%) | 32 (14.6%) | - | 16 (11.4%) | - |
| *Clinical Motor Features* |  |  |  |  |
| UPDRS-Part II | 5.6 (3.9) | 11.7 (7.6) | 5.9 (4.2) | 10.5 (6.7) |
| UPDRS-Part III | 21.3 (9.0) | 29.4 (14.3) | 19.0 (7.5) | 28.6 (11.9) |
| Schwab & England score | 94.4 (5.8) | 86.2 (12.7) | 92.6 (6.1) | 81.6 (12.9) |
| PIGD score | 0.2 (0.2) | 0.5 (0.6) | 0.2 (0.2) | 0.5 (0.5) |
| *Clinical Non-Motor Features* |  |  |  |  |
| UPDRS-Part I | 5.6 (4.1) | 7.8 (5.5) | 5.8 (3.7) | 8.5 (5.7) |
| Epworth sleepiness score | 5.9 (3.6) | 8.0 (5.2) | 5.5 (3.4) | 7.0 (4.2) |
| Geriatric depression scale (GDS) | 2.2 (2.5) | 3.0 (3.2) | 2.3 (2.0) | 3.0 (2.9) |
| State-trait anxiety inventory (STAI) score | 64.0 (18.3) | 66.1 (21.5) | 66.9 (18.1) | 66.6 (19.8) |
| Impulse control disorders (QUIP score) | 0.3 (0.8) | 0.4 (0.7) | 0.3 (0.9) | 0.4 (0.8) |
| REM sleep behavior disorder (RBD) score | 4.1 (2.6) | 4.9 (3.2) | 4.2 (2.7) | 5.1 (3.2) |
| SCOPA-AUT score | 9.3 (6.0) | 12.9 (7.3) | 9.7 (6.3) | 13.9 (7.7) |
| MOCA score | 27.3 (2.2) | 26.2 (3.9) | 26.7 (2.5) | 25.9 (4.1) |

*UPDRS: Unified Parkinson's disease rating scale; PIGD: postural instability and gait disturbance; SCOPA-AUT: Scales for Outcomes in PD-Autonomic (SCOPA-AUT); MOCA: Montreal Cognitive Assessment*

### Longitudinal Progression and Trajectories of Clinical Features

The baseline PD-network atrophy score showed significant association with all measures of progression at 54 (±12) months, including the GCO (r= −0.25, *p*=0.0002, **Figure 2**), motor dysfunction on UPDRS-parts II-III (r= −0.20, *p*=0.0027), PIGD score (r= −0.24, *p*=0.0004), non-motor manifestations on UPDRS-part I (r= −0.21, *p*=0.0014) and MoCA (r= 0.25, *p*=0.0002) (**Table 2**).

**Table 2.** Linear correlation coefficients for the associations between selected markers at baseline and changes in the outcomes of interest after follow-up (4.5 years). Data are presented as correlation coefficient (p-value).

|  | Δ MOCA Score | Δ UPDRS Non-Motor Score | Δ PIGD Score | Δ UPDRS Motor Score | Δ Schwab & England ADL Score | Δ Global Composite Outcome |
| --- | --- | --- | --- | --- | --- | --- |
| PD- atrophy score | **0.25**<br>**(0.0002)** | -0.16<br>(0.0189) | **-0.24**<br>**(0.0003)** | -0.18<br>(0.0064) | 0.06<br>(0.3652) | **-0.22**<br>**(0.0011)** |
| SBR (caudate) | 0.20<br>(0.0035) | -0.14<br>(0.0330) | -0.08<br>(0.2042) | -0.06<br>(0.3538) | 0.18<br>(0.0079) | -0.19<br>(0.0040) |
| SBR (putamen) | 0.12<br>(0.0794) | -0.08<br>(0.2160) | -0.05<br>(0.4377) | -0.01<br>(0.8993) | 0.07<br>(0.2663) | -0.09<br>(0.1936) |
| UPDRS-Motor (II-III) | -0.13<br>(0.0095) | -0.02<br>(0.7241) | 0.04<br>(0.4171) | **-0.22**<br>**(0.0000)** | 0.03<br>(0.5232) | -0.12<br>(0.0162) |
| PIGD score | -0.06<br>(0.2395) | -0.08<br>(0.0920) | **-0.25**<br>**(0.0000)** | -0.06<br>(0.2527) | -0.07<br>(0.1485) | -0.02<br>(0.7357) |
| CSF Aβ 42/T-tau ratio | 0.15<br>(0.0042) | -0.04<br>(0.4393) | -0.08<br>(0.1057) | -0.12<br>(0.0151) | 0.14<br>(0.0054) | 0.13<br>(0.0093) |
| CSF α-synuclein | 0.07<br>(0.1894) | -0.02<br>(0.6200) | -0.02<br>(0.6322) | -0.06<br>(0.2024) | 0.02<br>(0.7209) | -0.07<br>(0.1853) |

grey = not significant (p-value > 0.05)
Black = uncorrected significant (p-value < 0.05)
**Bold Blue = significant after Bonferroni correction p-value < 0.05 after bonferroni correction)**
**UPDRS:** Unified Parkinson's disease rating scale; **PD:** Parkinson's disease; **PIGD:** postural instability and gait disturbance; **SCOPA-AUT:** Scales for Outcomes in PD-Autonomic (SCOPA-AUT); **MOCA:** Montreal Cognitive Assessment; **CSF:** cerebrospinal fluid; **SBR:** striatum binding ratio

**Figure 2.**
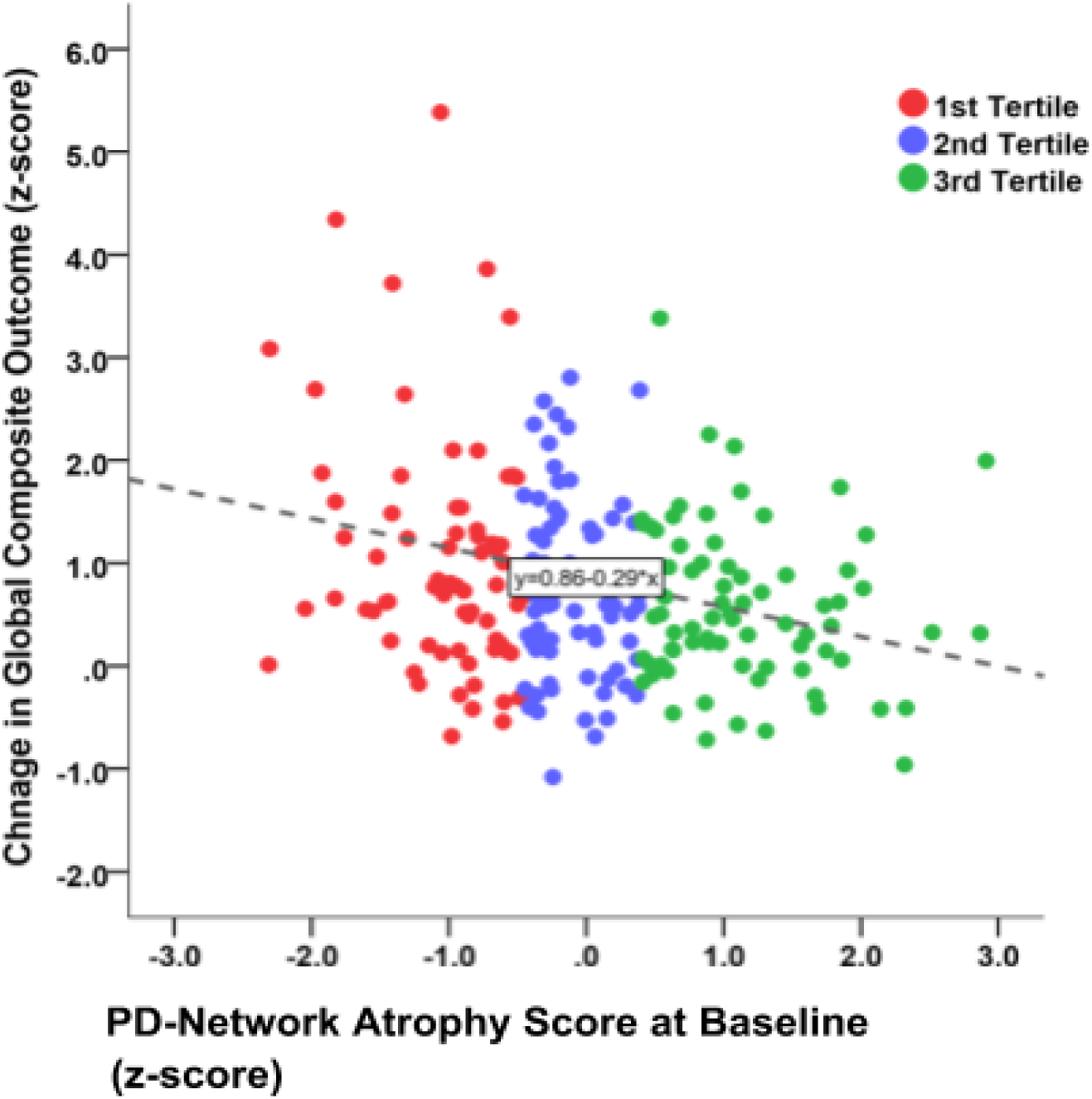
Association between the Parkinson’s disease (PD)-network atrophy score at baseline and progression rate in global composite outcome after an average of 4.5 years in people with PD (B= −0.240, p<0.001). Tertiles refer to the mean degree of atrophy.

When divided into tertiles based on the MRI-derived atrophy score (**Table 3, Figure 3**) those in the 1^st^ tertile (i.e. most atrophy) progressed significantly faster on motor signs, with a 3-unit further increase in UPDRS-part III each year (*p*=0.012) compared to the 3^rd^ tertile. They also experienced faster worsening of gait and postural instability (PIGD score progression=+0.21 (*p*<0.0001) compared to the 3^rd^ tertile). Subjects in the worst tertile not only had a lower MoCA score at baseline (−1.02, *p*=0.0089), but also declined faster by −0.74 (*p*=0.0372) extra MoCA points per year. Members of the 1^st^ tertile also demonstrated a+0.38 (*p*=0.0029) and +0.29 (*p*=0.0220) further annual increase in the GCO z-score compared to the 3^rd^ and 2^nd^ tertiles, respectively.

**Table 3.** Longitudinal trajectories of various clinical outcomes compared between tertiles of baseline of proposed biomarkers and the given outcome in the main dataset (3T MRI) and validation dataset (1.5T MRI) of PPMI population using mixed effect models. The first column represents model variables for each the proposed biomarker (separated in blue) and each column in this table shows an outcome. (e.g. first column first subsection represents the PD-ICA as a proposed biomarker and the outcome is based on UPDRS-III measured longitudinally in 3T MRI dataset. The corresponding model therefore will be UPDRS-part III ∼ 1 + Age + Sex + Education + Group + Age*Group + (1❘Subject) Group = Biomarker tertile, in this case Group = PD-ICA MRI score tertile As seen in the table the model has 61% variance explained (adjusted R-squared), There is a significant effect of age on UPDRS-III as well as a significant difference in UPDRS-III between most atrophied and least atrophied group after accounting for age, sex, and education. Furthermore, there is a significant interaction between the atrophy group and age suggesting for each extra year in age most atrophied group will gain three point more than least atrophied group in their UPDRS-III score

| dataset | 3T MRI (Main dataset) |  |  |  | 1.5T MRI (Validation dataset) |  |  |  |
| --- | --- | --- | --- | --- | --- | --- | --- | --- |
| outcome | UPDRS-III | Cognition | GCO | PIGD | UPDRS-III | Cognition | GCO | PIGD |
| PD-ICA | UPDRS-III | Cognition | GCO | PIGD | UPDRS-III | Cognition | GCO | PIGD |
| Age Effect | **5.20**<br>(3.24,7.16) | **-2.54**<br>(-4.50,-0.58) | **5.77**<br>(3.81,7.73) | **2.82**<br>(0.86,4.78) | **5.12**<br>(3.16,7.08) | -0.78<br>(-2.74,1.19) | **7.11**<br>(5.15,9.07) | **2.55**<br>(0.59,4.52) |
| Sex Effect (male) | 1.08<br>(-0.88,3.04) | **-2.25**<br>(-4.21,-0.28) | 1.59<br>(-0.37,3.56) | -0.28<br>(-2.24,1.68) | 1.02<br>(-0.95,2.98) | -0.61<br>(-2.57,1.36) | 0.58<br>(-1.39,2.54) | -0.46<br>(-2.42,1.50) |
| Education Effect | -0.10<br>(-2.06,1.86) | 2.07<br>(0.10,4.03) | -1.06<br>(-3.02,0.90) | -0.78<br>(-2.74,1.18) | 1.03<br>(-0.93,2.99) | **2.46**<br>(0.50,4.42) | -0.14<br>(-2.10,1.82) | 0.90<br>(-1.06,2.86) |
| Baseline Difference<br>2<sup>nd</sup> - 1<sup>st</sup> Tertile | -0.18<br>(-2.15,1.78) | 0.72<br>(-1.25,2.68) | -0.78<br>(-2.74,1.18) | -2.12<br>(-4.08,-0.16) | -1.75<br>(-3.71,0.21) | 1.15<br>(-0.82,3.11) | -3.05<br>(-5.01,-1.09) | **-2.52**<br>**(-4.48,-0.55)** |
| 3<sup>rd</sup> - 1<sup>st</sup> Tertile | **-2.59**<br>**(-4.55,-0.63)** | 1.75<br>(-0.22,3.71) | -3.06<br>(-5.03,-1.10) | -4.37<br>(-6.33,-2.41) | -3.70<br>(-5.66,-1.74) | 2.15<br>(0.19,4.12) | -5.70<br>(-7.66,-3.74) | **-5.65**<br>**(-7.61,-3.69)** |
| Age * Group Interaction<br>2<sup>nd</sup> - 1<sup>st</sup> Tertile | 0.17<br>(-1.79,2.13) | -1.10<br>(-3.06,0.87) | 0.74<br>(-1.22,2.70) | **2.23**<br>**(0.27,4.19)** | 1.35<br>(-0.61,3.31) | -1.30<br>(-3.26,0.66) | **2.62**<br>**(0.66,4.58)** | 2.54<br>(0.58,4.50) |
| 3<sup>rd</sup> - 1<sup>st</sup> Tertile | **2.52**<br>**(0.56,4.48)** | **-2.17**<br>**(-4.13,-0.20)** | **3.03**<br>**(1.06,4.99)** | **4.43**<br>**(2.47,6.39)** | **3.51**<br>**(1.55,5.47)** | **-2.36**<br>**(-4.33,-0.40)** | **5.01**<br>**(3.05,6.97)** | 5.49<br>(3.53,7.45) |
| **SPECT SBR** | UPDRS-III | Cognition | GCO | PIGD | UPDRS-III | Cognition | GCO | PIGD |
| Age Effect | **7.37**<br>**(5.41,9.33)** | **-5.14**<br>**(-7.11,-3.18)** | **6.61**<br>**(4.65,8.57)** | **4.12**<br>**(2.16,6.08)** | **7.23**<br>**(5.27,9.19)** | -0.22<br>(-2.18,1.75) | **8.13**<br>**(6.17,10.10)** | **4.56**<br>**(2.60,6.53)** |
| Sex Effect (male) | -0.37<br>(-2.33,1.59) | -1.01<br>(-2.97,0.96) | 0.59<br>(-1.37,2.55) | 0.55<br>(-1.41,2.51) | -0.76<br>(-2.72,1.20) | 0.57<br>(-1.39,2.54) | -1.27<br>(-3.24,0.69) | -1.70<br>(-3.66,0.27) |
| Education Effect | 1.68<br>(-0.29,3.64) | 0.38<br>(-1.58,2.34) | 1.74<br>(-0.23,3.70) | 0.52<br>(-1.44,2.48) | -0.32<br>(-2.28,1.64) | 1.60<br>(-0.36,3.57) | -1.25<br>(-3.21,0.71) | -1.04<br>(-3.01,0.92) |
| Baseline Difference<br>2<sup>nd</sup> - 1<sup>st</sup> Tertile | 2.15<br>(0.19,4.11) | -1.13<br>(-3.09,0.83) | 0.14<br>(-1.82,2.10) | -0.33<br>(-2.29,1.63) | 0.33<br>(-1.63,2.29) | 1.99<br>(0.02,3.95) | -1.55<br>(-3.51,0.41) | -1.91<br>(-3.88,0.05) |
| 3<sup>rd</sup> - 1<sup>st</sup> Tertile | 0.13<br>(-1.84,2.09) | 0.69<br>(-1.27,2.65) | -3.11<br>(-5.08,-1.15) | -3.57<br>(-5.54,-1.61) | 0.79<br>(-1.17,2.75) | 2.48<br>(0.51,4.44) | -1.65<br>(-3.62,0.31) | 0.01<br>(-1.95,1.97) |
| Age * Group Interaction<br>2<sup>nd</sup> - 1<sup>st</sup> Tertile | **-2.01**<br>**(-3.97,-0.04)** | 1.20<br>(-0.76,3.17) | -0.20<br>(-2.16,1.76) | 0.48<br>(-1.48,2.44) | -0.28<br>(-2.25,1.68) | **-2.35**<br>**(-4.31,-0.39)** | 1.43<br>(-0.53,3.39) | 1.77<br>(-0.19,3.73) |
| 3<sup>rd</sup> - 1<sup>st</sup> Tertile | 0.30<br>(-1.66,2.26) | -0.75<br>(-2.71,1.21) | **3.49**<br>**(1.53,5.45)** | **4.06**<br>**(2.10,6.02)** | -0.71<br>(-2.68,1.25) | **-2.69**<br>**(-4.66,-0.73)** | 1.34<br>(-0.62,3.31) | -0.08<br>(-2.04,1.88) |
| **PIGD** | UPDRS-III | Cognition | GCO | PIGD | UPDRS-III | Cognition | GCO | PIGD |
| Age Effect | **8.28**<br>**(6.32,10.24)** | **-5.27**<br>**(-7.23,-3.31)** | **8.89**<br>**(6.92,10.85)** | **6.12**<br>**(4.16,8.08)** | **6.63**<br>**(4.67,8.60)** | **-1.28**<br>**(-3.24,0.68)** | **8.17**<br>**(6.21,10.13)** | **4.05**<br>**(2.09,6.01)** |
| Sex Effect (male) | 0.12<br>(-1.84,2.08) | -0.44<br>(-2.40,1.52) | 0.24<br>(-1.73,2.20) | -0.97<br>(-2.93,0.99) | 0.90<br>(-1.06,2.86) | -0.58<br>(-2.54,1.39) | -0.18<br>(-2.14,1.78) | -0.50<br>(-2.47,1.46) |
| Education Effect | **3.15**<br>**(1.19,5.11)** | 0.02<br>(-1.94,1.99) | **2.27**<br>**(0.30,4.23)** | 1.72<br>(-0.24,3.68) | 0.68<br>(-1.28,2.64) | **2.51**<br>**(0.54,4.47)** | -0.81<br>(-2.77,1.15) | 0.22<br>(-1.74,2.18) |
| Baseline Difference<br>2<sup>nd</sup> - 1<sup>st</sup> Tertile | 1.25<br>(-0.71,3.21) | -0.60<br>(-2.56,1.36) | 1.55<br>(-0.41,3.52) | 0.93<br>(-1.03,2.89) | -1.44<br>(-3.40,0.52) | 1.23<br>(-0.73,3.19) | -1.19<br>(-3.15,0.77) | -0.88<br>(-2.85,1.08) |
| 3<sup>rd</sup> - 1<sup>st</sup> Tertile | **2.69**<br>**(0.73,4.65)** | -0.35<br>(-2.31,1.61) | 0.96<br>(-1.00,2.93) | 0.78<br>(-1.18,2.74) | 1.74<br>(-0.22,3.70) | 0.60<br>(-1.36,2.57) | -0.18<br>(-2.14,1.78) | 0.85<br>(-1.11,2.81) |
| Age * Group Interaction |  |  |  |  |  |  |  |  |
| 2<sup>nd</sup> - 1<sup>st</sup> Tertile | -0.79<br>(-2.75,1.17) | 0.39<br>(-1.57,2.35) | -1.22<br>(-3.18,0.74) | -0.38<br>(-2.34,1.58) | 1.79<br>(-0.18,3.75) | -1.40<br>(-3.36,0.57) | 1.45<br>(-0.51,3.42) | 1.38<br>(-0.58,3.34) |
| 3<sup>rd</sup> - 1<sup>st</sup> Tertile | **-2.20**<br>**(-4.16,-0.24)** | 0.10<br>(-1.87,2.06) | -0.31<br>(-2.27,1.65) | 0.34<br>(-1.62,2.30) | -1.68<br>(-3.64,0.29) | -0.92<br>(-2.88,1.04) | 0.36<br>(-1.61,2.32) | -0.17<br>(-2.13,1.79) |
| Alpha-synuclein | UPDRS-III | Cognition | GCO | PIGD | UPDRS-III | Cognition | GCO | PIGD |
| Age Effect | **5.02**<br>**(3.06,6.98)** | **-4.85**<br>**(-6.81,-2.88)** | **7.79**<br>**(5.82,9.75)** | **6.09**<br>**(4.13,8.06)** | **6.72**<br>**(4.76,8.68)** | **-3.79**<br>**(-5.75,-1.82)** | **8.52**<br>**(6.56,10.48)** | **6.51**<br>**(4.55,8.47)** |
| Sex Effect (male) | 1.49<br>(-0.47,3.46) | -1.29<br>(-3.25,0.67) | 1.89<br>(-0.07,3.85) | 0.11<br>(-1.85,2.07) | 0.36<br>(-1.60,2.32) | -0.43<br>(-2.39,1.53) | 0.88<br>(-1.08,2.84) | 1.06<br>(-0.90,3.02) |
| Education Effect | 1.45<br>(-0.51,3.41) | 1.52<br>(-0.44,3.48) | 0.63<br>(-1.33,2.59) | 0.67<br>(-1.29,2.63) | 1.14<br>(-0.82,3.10) | 0.62<br>(-1.35,2.58) | -0.44<br>(-2.40,1.52) | -0.84<br>(-2.80,1.12) |
| Baseline Difference |  |  |  |  |  |  |  |  |
| 2<sup>nd</sup> - 1<sup>st</sup> Tertile | -1.80<br>(-3.76,0.16) | -0.74<br>(-2.70,1.22) | -0.20<br>(-2.16,1.76) | 0.44<br>(-1.52,2.40) | -1.36<br>(-3.32,0.60) | -0.87<br>(-2.83,1.09) | -1.76<br>(-3.72,0.20) | 0.31<br>(-1.65,2.27) |
| 3<sup>rd</sup> - 1<sup>st</sup> Tertile | **-2.01**<br>**(-3.97,-0.05)** | 0.33<br>(-1.63,2.29) | -0.36<br>(-2.32,1.60) | -0.72<br>(-2.68,1.24) | 0.87<br>(-1.09,2.83) | **-2.03**<br>**(-3.99,-0.06)** | -0.33<br>(-2.30,1.63) | 1.86<br>(-0.10,3.82) |
| Age * Group Interaction |  |  |  |  |  |  |  |  |
| 2<sup>nd</sup> - 1<sup>st</sup> Tertile | 1.87<br>(-0.09,3.83) | 0.80<br>(-1.16,2.76) | 0.04<br>(-1.93,2.00) | -0.55<br>(-2.51,1.42) | 1.75<br>(-0.21,3.71) | 0.91<br>(-1.05,2.87) | 2.08<br>(0.12,4.04) | -0.33<br>(-2.29,1.63) |
| 3<sup>rd</sup> - 1<sup>st</sup> Tertile | 1.70<br>(-0.26,3.67) | -0.33<br>(-2.29,1.64) | -0.08<br>(-2.04,1.88) | 0.38<br>(-1.58,2.34) | -1.06<br>(-3.02,0.90) | 1.95<br>(-0.01,3.92) | 0.02<br>(-1.94,1.99) | **-2.29**<br>**(-4.25,-0.33)** |

UPDRS: Unified Parkinson's disease rating scale; PD: Parkinson's disease; ICA: independent component analysis; PIGD: postural instability and gait disturbance; MOCA: Montreal Cognitive Assessment; GCO: global composite outcome; SBR: striatal binding ratio
Each column in this table shows an outcome and each of the three subsections within a column represents one mixed effect model (e.g. first column first subsection represents UPDRS-part III ~ AGE + SEX + GROUP (PD-ICA MRI biomarker tertiles) + AGE\*GROUP)
\* p<0.00001, % p<0.0005, & p<0.001, ^ p<0.005, \$ p<0.01, # p<0.05

**Figure 3.**
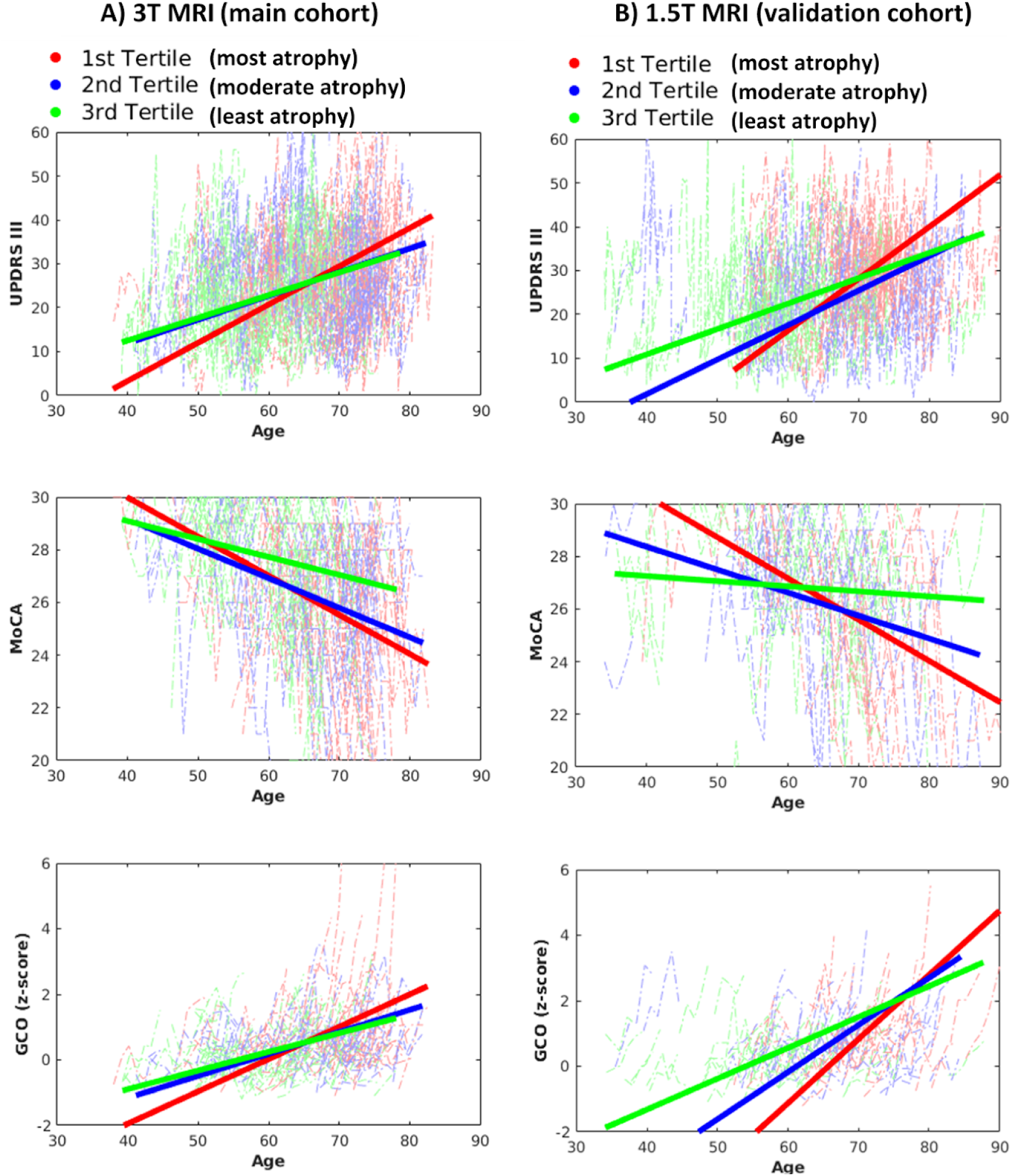
Longitudinal trajectories of the outcomes (i.e. motor UPDRS-part III, MoCA cognition, Global composite outcome scores) in different tertiles of the Parkinson’s disease (PD)-network atrophy score at baseline in 3T and 1.5T PPMI sub-populations (Mean follow-up duration in the entire population= 4.5 years)

### Comparison with Other Potential Biomarkers

We compared the predictive value of the MRI-derived atrophy measure to other potential biomarkers. ROC curve analysis showed that, among all biomarkers evaluated, only the baseline PD-network atrophy score significantly predicted a more than 1.5 SD worsening in GCO after 4.5 years of follow-up (AUC=0.64, *p*=0.005, **Figure 4**). Baseline PIGD score, motor severity (UPDRS-part II and III), and SPECT SBR in caudate and putamen either lacked sensitivity or statistical power to predict a 1.5 SD increase in GCO after this period (**Figure 4**). The baseline PIGD score failed to associate with longitudinal progression at 4.5 years. Most of the associations with the baseline CSF A*β*42/T-tau ratio disappeared after regressing out the effect of age. Compared to SPECT SBR, MRI PD-network atrophy score not only had larger regression coefficients but also survived Bonferroni correction for multiple comparisons (**Table 2**).

**Figure 4.**
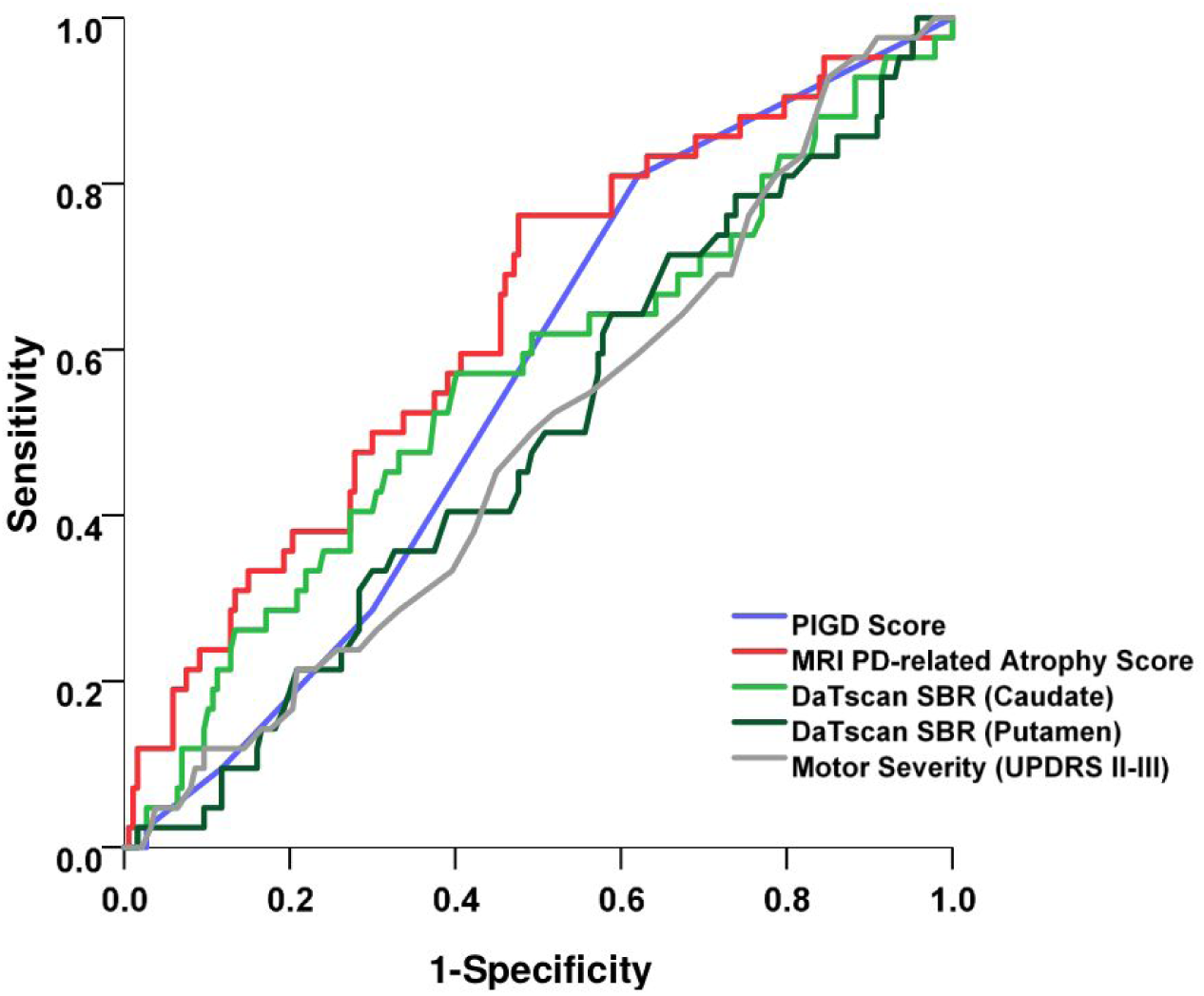
Receiver operating characteristics (ROC) analysis to compare the area under the curve (AUC) of baseline Parkinson’s disease (PD)-network atrophy score (AUC=0.639±0.047, p=0.005), UPDRS-Parts II and III (AUC=0.493±0.048, p=0.894), PIGD score (AUC=0.559±0.044, p=0.230), striatal binding ratio in caudate (AUC=0.560±0.051, p=0.224) and putamen (AUC=0.494±0.048, p=0.904) to predict 1.5 standard deviation (SD) increase in the global composite outcome after an average of 4.5 years of follow-up

As summarized in **Table 3**, we repeated the same mixed effect models by grouping the population into tertiles at baseline based on the other potential biomarkers in the PPMI sub-population with available 3-T MRI (n=222). Members of the 1^st^ tertile of SBR (i.e. the most severe dopaminergic denervation) had further annual disease progression by +2.57 (p=0.0409) increase in UPDRS-part III progression (only 1^st^ tertile vs 2^nd^ tertile), +0.43 (*p*=0.0003) increase in GCO z-score progression, and +0.16 (p=0.0003) increase in PIGD progression. The 3^rd^ tertile of PIGD baseline score (i.e. the most severe postural and gait disturbances), progressed the least in UPDRS-part III [-2.59 (p=0.0364) compared to the 1^st^ tertile], representing a ceiling effect for the progression of PIGD score after 4.5 years of initial diagnosis. While this effect is marginally significant, it shows lack of prognostic power of the baseline PIGD score. Neither the SPECT tertiles nor the PIGD motor phenotypes significantly predicted cognitive decline at follow-up. Finally, the MRI network atrophy score was a stronger predictor of rate of change in UPDRS-part III (R^2^=0.197, p<0.003) (and all other measures) than the UPDRS-part III itself (R^2^=0.122, p=0.070).

### Phenotype Shifting

Using previously published clinical criteria for subtyping PD,^18^ 118 participants were categorized as “*mild motor-predominant*” PD at baseline, of whom 72.9% remained in the same subtype after 4.5 years of follow-up. As illustrated in **Figure 5-A**, among those who were initially subtyped as “*mild motor-predominant*” PD, baseline PD-network atrophy was significantly worse in the ones who later shifted to the “*diffuse malignant*” subtype [-0.69 (SD=0.90) vs. 0.31 (SD=1.08), *p*=0.042]. A similar effect could also be seen for the “intermediate” subtype, where the baseline PD-network atrophy score was significantly lower in the subgroup who later shifted into the “*diffuse malignant*” subtype compared to those who shifted to “*mild motor-predominant*” [-0.60 (SD=1.03) vs. −0.01 (SD=0.91), *p*=0.034] (**Figure 5-A**).

**Figure 5.**
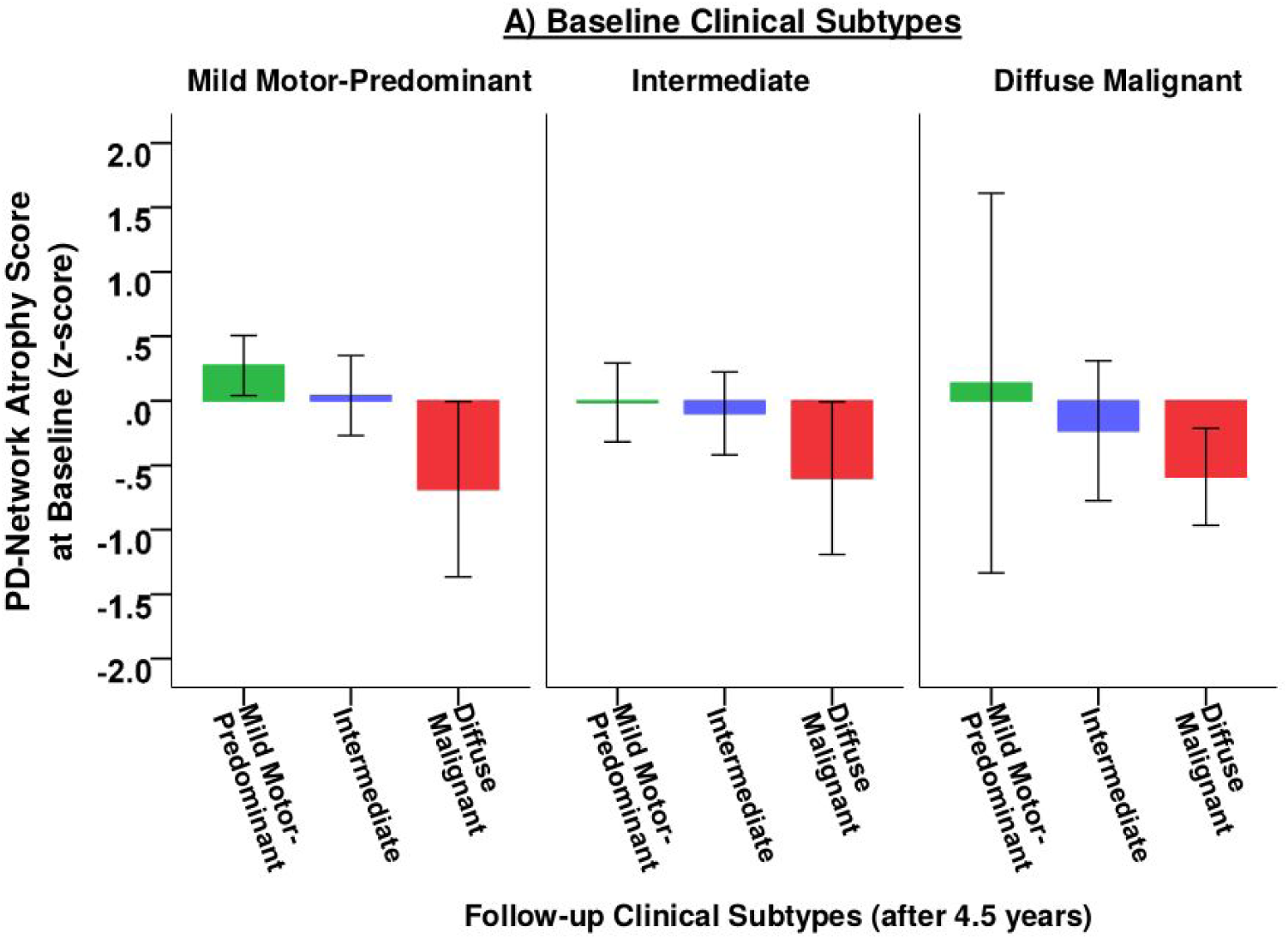
Prediction of the change in phenotypic subgroup assignment. Baseline value of Parkinson’s disease (PD)-network atrophy score at baseline among PPMI population categorized in different clinical subtypes at baseline and after 4.5 years of follow-up.

Furthermore, there was a significant difference in baseline PD-network atrophy score between individuals classified as stable “*mild motor-predominant*” and “*diffuse malignant*”subtypes after 4.5 years [0.31 (SD=1.08) vs. −0.59 (SD=0.62) vs., *p*=0.008]. Among individuals initially categorized in the tremor-dominant phenotype, baseline PD-network atrophy score was significantly lower in individuals who later progressed into the PIGD-dominant phenotype after 4.5 years, compared to those who remained tremor-dominant [-0.21 (SD=0.93) vs. 0.18 (SD=1.00), *p*=0.042].

### Longitudinal Progression and Trajectories of PD-network atrophy

To investigate the brain deformation alterations over time, we repeated our baseline neuroimaging analysis for the subsequent visits of PD and HC subjects. Here we used the PD-network atrophy score for each visit and used a mixed effect model for the longitudinal analysis. We found a significant main effect of age (*β*=, t=-9.27, p<0.0001) as well as a significant main effect of cohort(*β*=, t=-2.44,p = 0.01), however there was no significant interaction between age and cohort (*β*=, t=1.27, p=0.2). While a lack of significance is not evidence of lack of effect, based on current results we find that over the first 4-5 years after diagnosis most of the measurable is already present at baseline, while subsequent MRI scans demonstrate roughly equal aging-related tissue loss in PD patients and HC (**Figure 6-A**). While at the average group level the further increase in PD-network atrophy score in PD is attributable to aging, we further investigated the rate of progression for the PD-network atrophy score in each of our tertiles based on the baseline atrophy score. We also found a significant effect of age (*β*=-0.03, t=−10.7, p<0.0001), but in this analysis we find a significant interaction between age and tertile group both between the 1st tertile (i.e. most atrophy) and 3rd tertile (i.e. least atrophy) (*β*=−0.01, t=−2.13, p=0.03) and also between 1st tertile and 2nd tertile (i.e. intermediate atrophy) (*β*=−0.01, t=−1.97, p=0.05) (**Figure 6-B**). In the model we included the baseline tertiles to control for average atrophy levels at baseline between groups and these results are derived based on the slope of progression. Thus, only the PD patients with the most severe atrophy at baseline show ongoing atrophy that progresses faster than normal aging.

**Figure 6.**
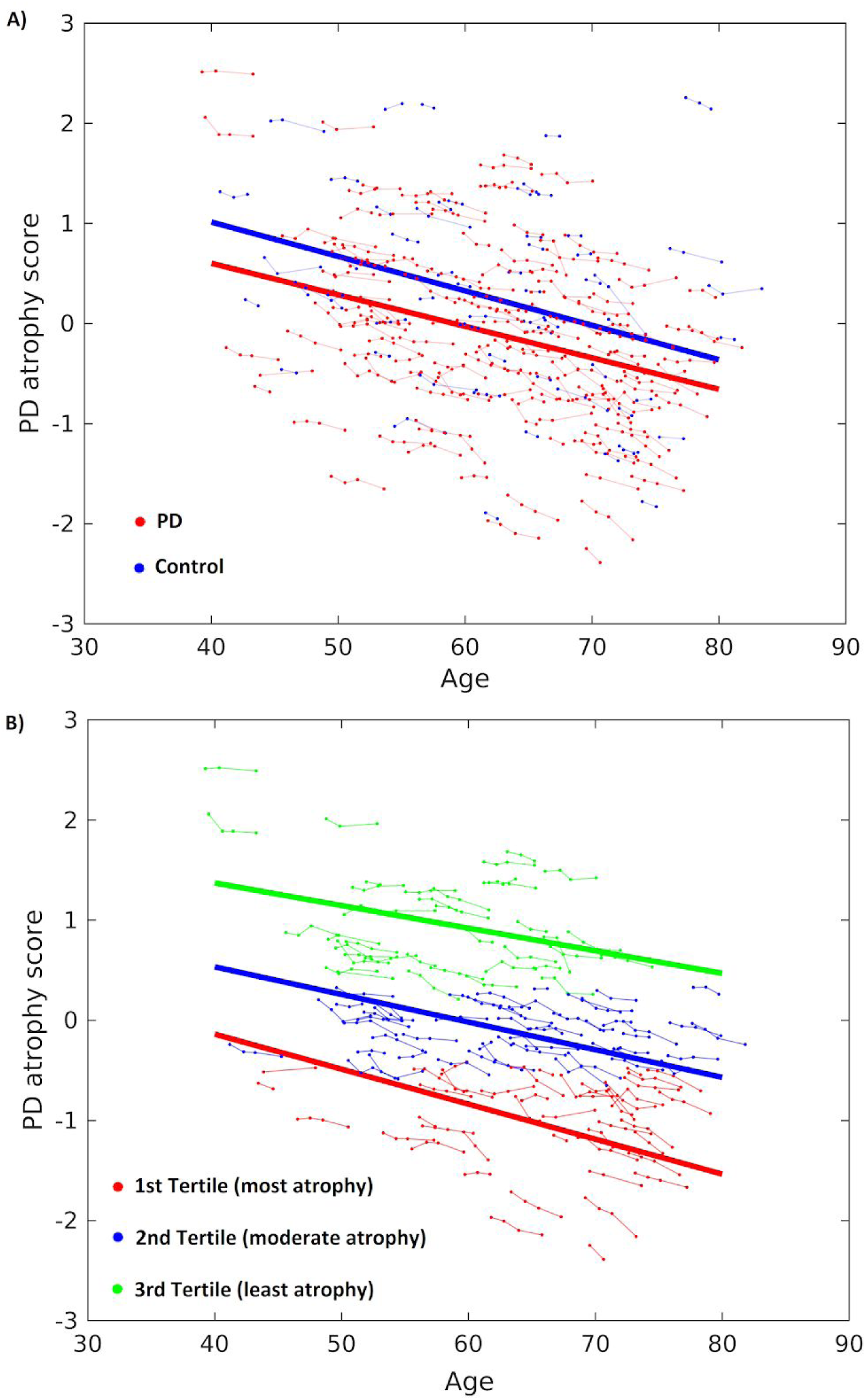
Longitudinal Progression and Trajectories of PD-network atrophy. A) The PD network atrophy score over time in healthy controls and PD patients. There is a significant main effect of cohort (PD - healthy controls) as well as a main effect of age. However, we didn’t find a significant interaction between cohort and age, suggesting that after the first hit in the PD-ICA network in PD patients, there is a dominant effect of aging in the following 3-4 years in both groups.B) While at the whole group level the dominant effect belongs to aging within PD patients, there is a significant interaction between age and disease severity (as measured by baseline tertiles in atrophy score). The patients with higher atrophy score at baseline show faster progression of the atrophy score in the following 3-4 years.

### Replication

To validate our findings, we repeated all mixed effect progression models in a separate sub-population of PPMI with 1.5-T MRI accessible at baseline (n=140, **Table 3**). All significant trajectory differences between the tertiles of PD-network atrophy score were replicated (**Figure 3**). Members of the 1^st^ tertile with the largest atrophy in the MRI-based PD-network at baseline, experienced a faster increase in UPDRS-part III (4-6 units/year), more rapid motor phenotype deterioration (0.21-0.36 PIGD scores/year), larger decline in MoCA (1-2 scores/year), and a worse overall prognosis on GCO (0.55-1.07 z-scores/year) after 4.5 years. By comparison, except for the cognitive domain where the 1^st^ tertile of the SBR showed faster decline compared to the 3^rd^ tertile, SBR subgroups and PIGD motor phenotype failed to show any significant difference in the trajectory of motor and global outcomes.

## DISCUSSION

This study has found that a single MRI-derived structural atrophy measure can be used as an early biomarker for PD prognosis. This PD-network atrophy measure combines atrophy in midbrain, basal ganglia, basal forebrain, medial temporal lobe, and discrete cortical regions into a single score for each patient. This score predicted the progression of motor impairment, clinical severity phenotype, cognition and a global measure of PD severity. All these results remain significant after regressing out the effect of age and were replicated in a validation analysis in an independent sample.

The PD-network atrophy score was a better predictor of prognosis than competing biomarkers. PIGD score, which has long been used as an indicator of motor phenotype and disease severity^21^ failed to predict progression in most motor and non-motor outcomes of interest at 4.5 years of follow-up. The MDS-UPDRS score at baseline was associated with cognitive decline, but showed no relation with other non-motor markers or even progression in the UPDRS score itself. CSF A*β*42/T-tau ratio was marginally associated with disease progression, however, this relationship failed to remain significant after regressing out the effect of age. SPECT imaging with ^123^| ioflupane, previously shown to correlate with cognitive impairment in *de novo* PD in this cohort^22^, had only modest predictive power for subsequent cognitive decline; moreover, our validation analysis demonstrated that it could not predict motor progression with the same strength as the MRI network atrophy score. The effect sizes of differences in trajectories were more prominent between the tertiles of the MRI network atrophy score for both motor and cognitive outcome measures, compared to SPECT.

It is also noteworthy that the baseline PD-network atrophy score could forecast a shift in PD subtype attribution after 4.5 years. Participants initially labelled as “*mild motor-predominant*” who later progressed into the “*diffuse malignant*” subtype had significantly larger atrophy in this MRI network at baseline compared to those with stable phenotype. The atrophy measure was also able to predict which patients in the “*intermediate*” subtype shifted to “*mild motor-predominant*” and “*diffuse malignant*” subtypes. This finding is particularly relevant for clinical practice as it might enable clinicians to anticipate future progression rate in newly diagnosed individuals who appear mildly affected early in disease, but will progress rapidly to more severe stages. Such a biomarker might also be useful for clinical trials of neuroprotective therapies, allowing the stratification of individuals predicted to progress more rapidly.

Other studies have suggested potential biomarkers for PD progression. These generally have targeted only one specific domain of PD (e.g. either motor or cognition) or lack clear clinical applicability at the individual level. Using machine-learning approaches on various clinical, imaging, CSF and genetic data, one recent study concluded that imaging data was unable to predict motor progression in *de novo* PD.^23^ However, in this study, DAT SPECT was the only imaging biomarker.^23^ We also found SPECT to be poor at predicting disease progression. On the contrary, another study using radiomics analysis on SPECT images at baseline and one year was able to predict changes in UPDRS-part III score in a small subset of patients from the PPMI cohort.24 There are several reasons why SPECT may be a better diagnostic than prognostic marker of PD: Parkinsonian medications may affect DAT SPECT uptake, there is likely a floor effect, and it is also possible that atrophy in the occipital lobe in PD may introduce noise in the measurement as it is the reference region typically used for measuring striatal DAT binding.

Other MRI biomarkers have been proposed. In one study, serial diffusion MRI measures of the free water in posterior substantia nigra were shown to correlate with the change in Hoehn and Yahr scale over 4 years.25 However, unlike these SPECT and MRI-based measures, the PD-network atrophy score proposed here describes a single measure at baseline that can predict disease progression in both motor and non-motor outcomes, and is easily obtainable using standard MRI pulse sequences.

A possible explanation for the prognostic strength of the MRI based biomarker is that it takes into account the entire spatial distribution of PD pathology. Although PD was initially characterized as a predominantly motor disorder caused by loss of dopamine neurons in the substantia nigra, several sources of evidence suggest this is an incomplete picture of the disease. Post-mortem analyses have suggested that PD is a spreading process that moves stereotypically from brainstem to subcortical regions to cortex.^26^ Clinical studies have demonstrated the importance and ubiquity of non-motor features such as autonomic, sleep, and cognitive dysfunction, attributable to widespread neuropathology below and above the substantia nigra.^27^ Our previous work with this PD network atrophy measure is supportive of a propagating process.^7,28,29,30^ Moreover, the discovery that neurotoxic misfolded *α*-synuclein can propagate trans-neuronally provides a mechanism for the propagation hypothesis first proposed by *Braak et al*.^31^ This suggests that the pattern of disease could be stereotyped in its spatial distribution as determined by the brains’ normal wiring diagram (or connectome)^7^. If so, an imaging measure that takes into account the entire distribution of atrophy could have more power to reflect global disease effects.

Some limitations of this study should be acknowledged. Follow-up duration of PPMI is still relatively short (4.5 years) and our findings need to be re-evaluated after a longer follow-up period. While our results have been validated in an independent sample from PPMI (a multi-center, multi-scanner dataset), future studies using external cohorts with different scanners and acquisition protocols can further investigate the generalizability of our proposed biomarker. The ROC analysis (**Figure 4**) shows that the PD-network biomarker outperforms the UPDRS, SPECT and PIGD scores as a prognostic indicator; however, even the PD-network score exhibits a significant trade-off between sensitivity and specificity. It’s ability to accurately predict outcomes in a single patient is limited; however, in the setting of a clinical trial, it remains a better prognostic marker than the others tested here.

Our study focused on baseline markers as predictors; future studies should also expand our analysis of the change in brain atrophy measures over time. We did however find that our mostly subcortical MRI atrophy network demonstrated progression in tissue loss that was no different than the effect seen in the age matched control group. Future studies should look for development and progression of atrophy in other brain areas. On the other hand, our study has some methodological strengths. By using mixed effect models, we analyzed all data points over time to increase statistical power, and adjusted for the effect of normal aging unlike most previous studies. We validated our analysis in an independent sample (PPMI population with 1.5T MRI), none of whose scans were used to develop the network atrophy score^7^. Finally, one of the advantages of our analysis was the simultaneous consideration of motor, cognition, and a multifaceted global outcome.

In conclusion, a PD-network atrophy measure based on a whole-brain MRI analysis can be more sensitive to disease severity and prognosis than more specific biomarkers that measure only dopaminergic deficit (e.g. DAT SPECT^24,25^) or MR-measures focused on the substantia nigra.^32^ Our biomarker’s ability to predict phenotype shifting highlights its clinical applicability, as well as the need to incorporate biomarker information in PD subtyping. The MRI analysis pipeline has been made freely accessible^7^ so that the corresponding atrophy score for any individual can be easily calculated for external applications. Further investigation to explore how different regional clusters of the PD-network atrophy might relate to the heterogenous pathology and clinical manifestations of PD is warranted.

## Acknowledgment

This work was funded by grants from the Michael J Fox Foundation for Parkinson’s Research, the Alzheimer’s Association, the Weston Brain Institute, the Canadian Institutes of Health Research, the Healthy Brains for Healthy Lives (HBHL) initiative of McGill University, the Preston Robb Fellowship and the Richard and Edith Strauss Scholarship (Faculty of Medicine, McGill University) in Canada. Data used in the preparation of this article were obtained from the Parkinson’s Progression Markers Initiative (PPMI) database (www.ppmi-info.org/data). For up-to-date information on the study, visit: www.ppmi-info.org. PPMI – a public-private partnership – is funded by the Michael J. Fox Foundation for Parkinson’s Research and funding partners, including AbbVie, Avid Radiopharmaceuticals, Biogen, Bristol-Myers Squibb, Covance, GE Healthcare, Genentech, GlaxoSmithKline (GSK), Eli Lilly and Company, Lundbeck, Merck, Meso Scale Discovery (MSD), Pfizer, Piramal Imaging, Roche, Sanofi Genzyme, Servier, Teva, and UCB (www.ppmi-info.org/fundingpartners).

